# Integrative cross tissue analysis of gene expression identifies novel type 2 diabetes genes

**DOI:** 10.1101/108134

**Authors:** Jason M. Torres, Alvaro N. Barbeira, Rodrigo Bonazzola, Andrew P. Morris, Kaanan P. Shah, Heather E. Wheeler, Graeme I. Bell, Nancy J. Cox, Hae Kyung Im

## Abstract

To understand the mechanistic underpinnings of type 2 diabetes (T2D) loci mapped through GWAS, we performed a tissue-specific gene association study in a cohort of over 100K individuals (*n_cases_* ≈ 26K, *n*_controls_ ≈ 84K) across 44 human tissues using MetaXcan, a summary statistics extension of PrediXcan. We found that 90 genes significantly (FDR < 0.05) associated with T2D, of which 24 are previously reported T2D genes, 29 are novel in established T2D loci, and 37 are novel genes in novel loci. Of these, 13 reported genes, 15 novel genes in known loci, and 6 genes in novel loci replicated (FDR_rep_ < 0.05) in an independent study (*n*_cases_ ≈ 10K, *n*_controls_ ≈ 62K). We also found enrichment of significant associations in expected tissues such as liver, pancreas, adipose, and muscle but also in tibial nerve, fibroblasts, and breast. Finally, we found that monogenic diabetes genes are enriched in T2D genes from our analysis suggesting that moderate alterations in monogenic (severe) diabetes genes may promote milder and later onset type 2 diabetes.

## Introduction

Type 2 diabetes (T2D) is a complex disease characterized by impaired glucose homeostasis resulting from dysfunction in insulin-secreting pancreatic islets and decreased insulin sensitivity in peripheral tissues [1]. In addition to environmental factors such as a sedentary lifestyle and poor diet, genetic susceptibility is an important contributor to the development of T2D [2]. Genome-wide association studies (GWAS) have uncovered more than 100 loci that significantly associate with either T2D or glucose-related traits [3, 4, 2]. However, the majority of single nucleotide polymorphisms (SNPs) significantly associated with T2D reside in intronic and intergenic regions rather than protein-encoding regions [5, 6]. The results from GWAS suggest an important role for genetic variation that regulates gene expression rather than altering codon sequence [7] and have motivated efforts to map the regulatory landscape of the genome [8, 9, 10]. Indeed, sets of trait-associated SNPs are enriched for variants that associate with gene expression (i.e. expression quantitative trait loci or eQTLs) [11] and that occupy DNAse hypersensitivity sites (DHS) [12] - regions overrepresented for eQTLs *per se* [13]. Moreover, DHS explain a disproportionately high share of SNP heritability [14] across 11 complex traits [15] and eQTLs mapped in insulin-responsive peripheral tissues similarly “concentrate” SNP heritability estimates for T2D [16].

Recent efforts to elucidate the functional consequences of non-coding disease-associated variants have challenged the assumption that the nearest gene to an associated marker is the relevant disease gene. For example, a non-coding SNP (rs12740374) within *CELSR2* at the 1p13 locus associated with myocardial infarction (MI) and low-density lipoprotein cholesterol (LDL-C) creates a C/EBP transcription factor binding site and alters the expression of *SORT1* (located ≈ 35 kb downstream of rs12740374) in primary human hepatocytes [17]. Moreover, *Sort1* knockdown and overexpression studies in mice altered LDL-C and very low density lipoprotein (VLDL) levels [17]. In a study of the *FTO* locus harboring the strongest association with obesity, researchers observed a long-range interaction between the associated intronic region of *FTO* and the promoter of *IRX3,* a downstream transcription factor located ≈ 500 kb away, but not with the *FTO* promoter [18]. Perturbing *IRX3* expression in the hypothalamus also reduced body mass accumulation in the background of a high fat diet and improved measures of metabolic health [18]. Obesity-associated SNPs within the locus were also significantly associated with *IRX3* expression in human cerebellum but not with *FTO* expression [18]. These examples demonstrate that regulatory consequences of disease-associated variants may not solely target the putative causal gene reported from GWAS, if at all. Thus, it is unclear to what extent regulatory genetic variation supports the putative causal gene at disease-associated loci. We sought to address this problem systematically by applying a statistical method that leverages the wealth of genotype and expression data from large-scale eQTL mapping studies.

Experimental techniques that manipulate endogenous gene expression (e.g. gene silencing, conditional knockout) can delineate relevant disease genes but are generally not suitable for *in vivo* human studies [19, 20]. By testing for association between the *genetic component* of gene expression and disease, we exploit the fact that nature essentially perturbs gene expression through random genetic variation introduced during meiosis. This analytic approach - implemented in the program PrediXcan - allows for a gene-based test that reflects the mechanism of transcription and presents advantages over GWAS and other study designs [21]. Namely, it reduces the multiple-testing burden, obviates causality issues encountered in differential gene expression studies, provides direction of effect for associated genes, and may implicate disease-relevant tissues [21]. Moreover, PrediXcan can corroborate reported disease genes as well as implicate novel genes as was the case for an analysis of type 1 diabetes based on predictors of gene expression in whole blood tissue [21]. In the present study, we applied a recent adaptation of the PrediXcan method - MetaXcan - that inputs summary GWAS data (Barbeira et al. 2016) to perform a systematic *in silico* evaluation of gene-level associations at T2D loci [22, 21]. We applied MetaXcan using predictive models corresponding to more than 40 human tissues to summary data from a trans-ethnic GWAS meta-analysis representing over 100K individuals and replicated results in an independent cohort.

## Results

### Genome-wide and cross-tissue scan of gene associations corroborates known T2D genes and implicates novel ones

We compared genetically regulated expression levels in T2D cases and controls from a trans-ethnic meta-analysis of GWAS (*n*_cases_ = 26, 488 and *n*_controls_ = 83, 964 from European, East Asian, South Asian, and Mexican American origin [23]) across 44 human tissues using reference transcriptome data. The differential expression of the genetic component was inferred using MetaXcan [22] with gene expression prediction models trained in RNAseq data from the Genotype-Tissue Expression Project (GTEx) [9]. We included an additional set of predictors trained in whole blood from the Depression Genes and Networks study where the available sample size (*n* = 922) is greater than that currently available for whole blood from GTEx (*n* = 338) [10].

Figure 1 shows a Manhattan plot of the full set of results across all genes and tissues (A) and qq-plots of the full set (B), the subset of genes within 1Mb of known T2D loci (C), and genes outside of known loci (D). Most of the significant genes are located in the vicinity of known T2D regions. After adjusting for the number of tests performed across all 44 tissue models (204,981 tests), we found 49 significant associations corresponding to 20 genes at the stringent Bonferroni threshold (*p* < 2.4 × 10^−7^) (See Table 1 and Supplementary Table S1). Of these 20 genes, 12 corresponded to those previously reported (nearest to the top T2D-associated SNP), 5 were novel but in the vicinity of known loci, and 3 were completely novel. When using FDR < 0.05, 90 genes were significantly associated with T2D risk; 22 of them were previously reported T2D genes, 31 were novel genes in established T2D loci, and 37 were novel genes in novel loci. (See Supplementary Table S2)

**Figure 1.**
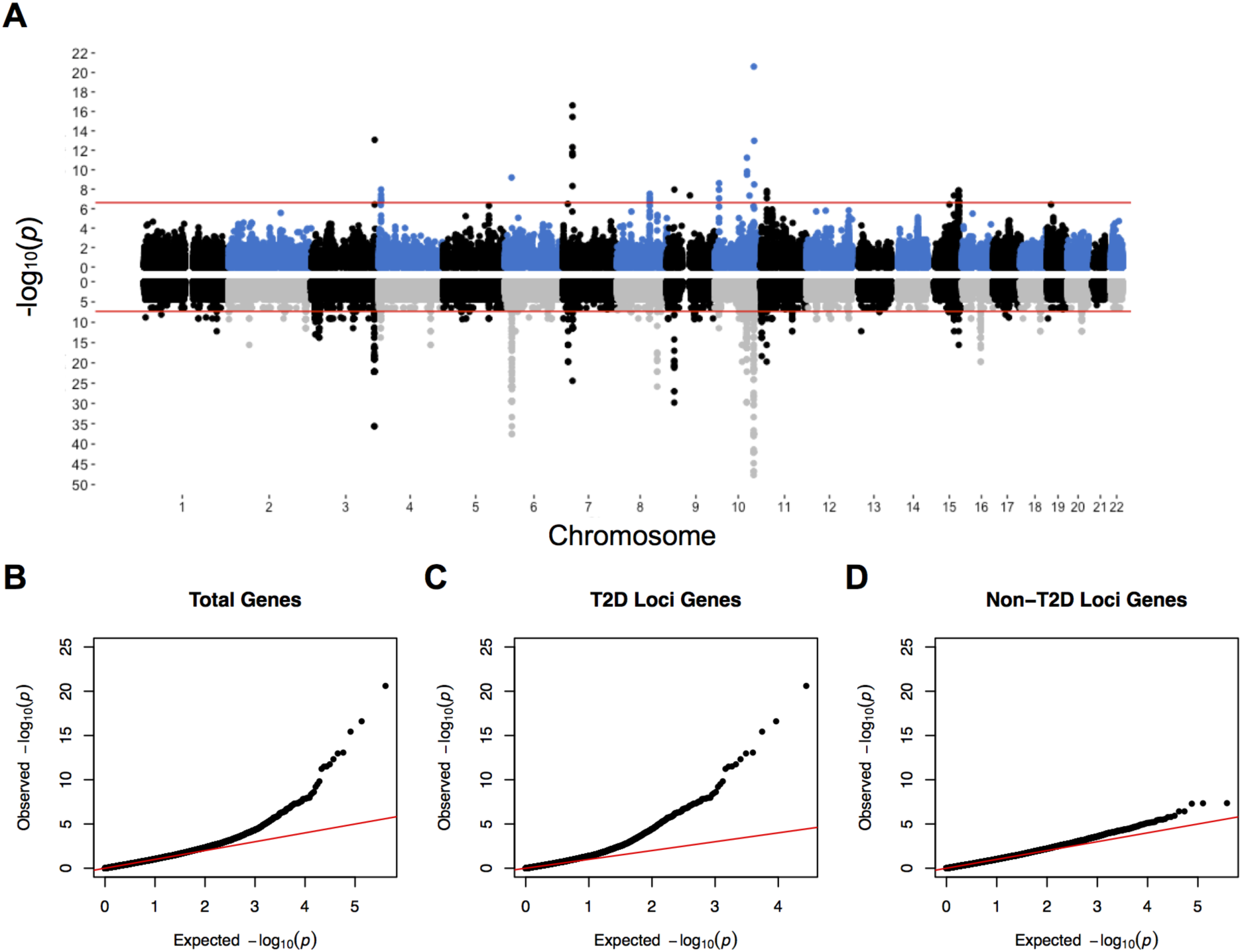
Tissue-level predicted gene expression associations map to predominantly known T2D loci. (A) (*Upper*) Manhattan plot showing MetaXcan gene associations across 44 tissue models using summary statistics from a trans-ethnic meta-analysis [23]. Red line denotes the Bonferroni significance threshold adjusted for the total number of tests performed across all tissue models (*p*< 2.4 10^−7^). Positions correspond to transcription start sites. (*Lower*) Manhattan plot showing SNP associations from the trans-ethnic meta-analysis of GWAS. Red line denotes marginal significance threshold (*p* =5 × 10^−6^). Y-axis is truncated at *−log*_10_(*p*) = 50 to enable comparison with MetaXcan profile as chromosome 10 association at *TCF7L2* locus would dominate plot. QQ-plot of tissue-level associations across 44 models for (B) total genes, (C) genes within 1 Mb of GWAS associations (*p* < 5 × 10^−6^), and (D) genes greater that 1 Mb away from GWAS associations.

**Table 1.** Significant association between predicted expression and T2D. Bonferroni corrected for all gene tissue pairs tested. When multiple tissues were significant, the top tissue result is shown.

| type | gene | chrom | reported.genes | disc.pval | gera.pval | gera.qval | meta.pval |
| --- | --- | --- | --- | --- | --- | --- | --- |
| T2D.Gene | AP3S2 | 15 | AP3S2,PRC1,VPS33B | 1.90E-07 | 4.60E-04 | 9.33E-04 | 2.20E-10 |
| T2D.Gene | CAMK1D | 10 | CDC123,CAMK1D | 2.40E-09 | 6.60E-04 | 1.16E-03 | 2.90E-10 |
| T2D.Gene | CCNE2 | 8 | DPY19L4,INTS8,CCNE2,<br>TP53INP1,NDUFAF6 | 3.00E-08 | 0.31 | 1.95E-01 | 3.00E-06 |
| T2D.Gene | HHEX | 10 | IDE,KIF11,HHEX | 5.90E-12 | 1.30E-04 | 5.17E-04 | 1.20E-12 |
| T2D.Gene | HMG20A | 15 | PEAK1,HMG20A,LINGO1 | 4.50E-08 | 6.50E-04 | 1.16E-03 | 6.70E-10 |
| T2D.Gene | IGF2BP2 | 3 | C3orf65,IGF2BP2 | 8.60E-14 | 4.80E-09 | 1.01E-07 | 1.10E-18 |
| T2D.Gene | JAZF1 | 7 | JAZF1 | 2.50E-17 | 2.40E-08 | 3.66E-07 | 5.90E-21 |
| T2D.Gene | NCR3LG1 | 11 | NCR3LG1,KCNJ11,ABCC8 | 1.50E-08 | 8.40E-05 | 4.52E-04 | 1.30E-10 |
| T2D.Gene | TCF7L2 | 10 | TCF7L2 | 2.50E-21 | 8.70E-19 | 7.95E-17 | 2.00E-31 |
| T2D.Gene | TP53INP1 | 8 | DPY19L4,INTS8,CCNE2,<br>TP53INP1,NDUFAF6 | 6.90E-08 | 0.57 | 3.12E-01 | 5.10E-06 |
| T2D.Gene | WFS1 | 4 | WFS1,PPP2R2C | 1.10E-08 | 4.70E-06 | 3.07E-05 | 4.30E-11 |
| Known.Region | CDKN2A | 9 | CDKN2B,DMRTA1 | 1.10E-08 | 0.003 | 3.61E-03 | 4.50E-10 |
| Known.Region | CYP26C1 | 10 | IDE,KIF11,HHEX | 1.50E-10 | 0.0053 | 5.84E-03 | 9.30E-10 |
| Known.Region | DCLRE1A | 10 | TCF7L2 | 1.10E-13 | 4.70E-07 | 4.77E-06 | 4.90E-17 |
| Known.Region | HLA-A | 6 | POUF5F1,TCF19 | 5.10E-08 | 0.5 | 2.80E-01 | 7.50E-06 |
| Known.Region | ID4 | 6 | CDKAL1 | 6.30E-10 | 2.40E-06 | 1.69E-05 | 2.10E-15 |
| Known.Region | NUDT5 | 10 | CDC123,CAMK1D | 1.10E-08 | 4.90E-04 | 9.33E-04 | 2.00E-09 |
| Known.Region | RCCD1 | 15 | AP3S2,PRC1,VPS33B | 1.40E-08 | 0.0018 | 2.53E-03 | 5.70E-10 |
| Known.Region | RCCD1 | 15 | AP3S2,PRC1,VPS33B | 1.40E-08 | 0.0012 | 1.92E-03 | 6.10E-10 |
| Unknown | ANKRD20A1 | 9 | none reported | 4.40E-08 | 0.36 | 2.18E-01 | 1.90E-06 |
| Unknown | CWF19L1 | 10 | none reported | 4.60E-08 | 0.45 | 2.56E-01 | 2.90E-06 |

The strongest gene association corresponds to *TCF7L2* - the gene harboring the strongest SNP-level association with T2D - and provides corroborating evidence that *TCF7L2* is the effector gene regulated by the non-coding variant driving the GWAS signal (Table S1 and Figure 1). This analysis provides additional *in silico* support for established T2D genes including *JAZF1, HHEX, WFS1, IGF2BP2,* and *CAMK1D.*

### Significant genes beyond known T2D loci: novel loci

Although most MetaXcan-implicated genes mapped to within 1 Mb of T2D-associated SNPs from the trans-ethnic meta-analysis of GWAS (*p* < 5 × 10^−6^), there were a few located beyond these intervals and hence designated as *genes in novel T2D loci*. Two associations met the stringent Bonferroni-corrected significance: *ANKRD20A1* and *CWF19L1* in breast mammary tissue (Table S1). Of the 90 genes implicated by associations at FDR ≤ 0.05, 37 mapped to *novel T2D loci* and included genes encoding potassium ion transporters (*KCNK17* and *KCNK7*) and zinc-finger proteins (*ZNF703, ZNF34*, and *ZNF771*) (Supplementary Table S3). Other *novel T2D loci* genes that were supported by two or more tissue-level associations include *MEIS1, JUND, MRPS33, TCP11L1, VIPAS39*, and *SNX11* (Supplementary Table S3). Collectively, these genes represent a class of discoveries that would have evaded detection in GWAS not only due to their distal location relative to significantly-associated marker SNPs but also due to proximal marker SNPs not meeting traditional genome-wide significance used for GWAS studies.

### Significant genes enriched in relevant pathways

To glean insight into relevant biological pathways, we performed a gene set enrichment analysis on the set of FDR ≤ 0.05 significant genes and found top Gene Ontology Biological Process (GO:BP) pathways to involve the insulin-secretory pancreatic *β*-cell (e.g. negative regulation of type B pancreatic cell apoptotic process, Supplemental Figure S4). This was also the case when we restricted this analysis to the set of *reported* T2D genes (Supplemental Table S5). However, we found fatty acid homeostasis to be a top pathway enriched among the set of *novel* T2D genes implicated in our MetaXcan analysis, underscoring a genetic contribution from variants regulating gene expression in insulin-responsive peripheral tissues (Supplemental Table S6).

### Enrichment of genes reported for related traits

We also explored shared etiology with other complex diseases by comparing our set of MetaXcan-implicated T2D genes with sets of genes implicated by GWAS listed in the NHGRI-EBI online catalogue. Unsurprisingly, we found that the set of MetaXcan-significant genes nearest to associated SNPs from the trans-ethnic study (i.e. *reported* T2D genes) were enriched among gene sets annotated to type 2 diabetes (*p* = 0.0001), fasting glucose-related traits with BMI interaction (*p* = 0.001), two-hour glucose challenge (*p =* 0.007), and glycated hemoglobin levels (*p =* 0.028) (Supplementary Table S7). However, an analysis based on the set of *novel* T2D genes (i.e. genes distal to associated SNPs from the trans-ethnic study at *p* < 5 × 10^−6^) revealed an enrichment for epilepsy (*p* = 0.0001) attributable to novel genes *COPZ2, SNX11,* and *MAST4* (Supplementary Table S8). Similarly, novel T2D genes *CCDC92, HOXA11, MEIS1,* and *JUND* were responsible for an observed enrichment for BMI-adjusted waist-to-hip ratio (*p* = 0.0004). *HKDC1* is a *novel* T2D gene implicated by our MetaXcan analysis that has been previously implicated in pregnancy-related glycemic traits and is the driver of the observed enrichment for this phenotype (*p* = 0.044) (Supplemental Table S8).

### Replication of novel T2D genes in independent GERA study

For the replication, we used 9,747 T2D cases and 61,857 controls from the Resource for Genetic Epidemiology Research on Adult Health and Aging study (GERA, phs000674.v1.p1). This independent dataset arises from a collaboration between the Kaiser Permanente Research Program on Genes, Environment, and Health and the UCSF Institute for Human Genetics represents a multi-ethnic cohort of 100K^+^ individuals from Northern California with available electronic medical records (EMRs). We performed MetaXcan analyses using GWAS results [24] and the same 44 human tissue expression models as in the discovery analysis.

Of the 90 top genes chosen for replication (discovery FDR < 0.05), 34 replicated (*p* < 0.05); 13 were previously reported genes, 15 were novel genes in known loci, and 6 were novel genes in novel loci. Moreover, the direction of effect was consistent across replicated associations (Figure 2).

**Figure 2.**
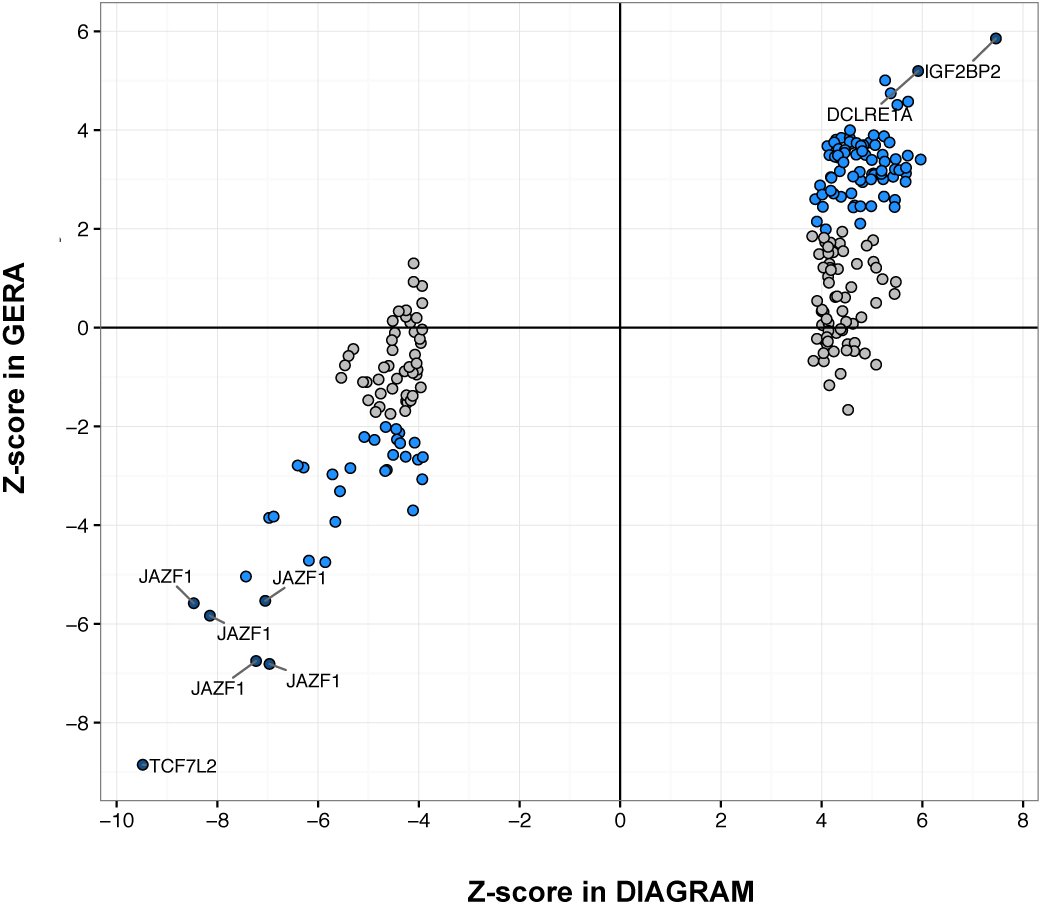
Replication of tissue-level gene associations in the GERA study. 106 of 207 tissue-level gene associations at FDR < 0.05 from the MetaXcan analysis of the trans-ethnic study meet the *p* < 0.05 threshold in the MetaXcan analysis of the GERA T2D study (blue). Gene associations meeting Bonferroni significance in GERA are labeled. All replicated associations show consistent direction of effect between studies.

Interestingly, decreased expression of *HKDC1* in aortic artery (*p* = 0.024) replicated in GERA. We observed replication for increased expression of *C2* in subcutaneous adipose tissue (*p* = 4.9 × 10^−4^), *HOXA11* in sigmoid colon (*p* = 0.0016), and *CYP21A2* in visceral (omentum) adipose tissue (*p* = 0.046). The remaining set of *novel* genes replicated at regions spanning T2D-associated loci (i.e. T2D windows) include *EVC, ID4, EXT1, NUDT5, CYP26C1, DCLRE1A, GPAM, NHLRC2, RCN2,* and *CTD-2021H9.3* (Supplementary Table S9).

Among reported genes that replicated in GERA are *JAZF1, HHEX, WFS1, CAMK1D, NCR3LG1, AP3S2, HMG20A, CDKN2A, KCNJ11, IRS1,* and IGF2BP2 (Table S1)

Five genes outside of known T2D regions replicated in GERA. These include *KCNK17* (potassium two pore domain channel subfamily K member 17), two zinc finger protein encoding genes, *ZNF703* (zinc finger protein 703) and *ZNF771* (zinc finger protein 771), *PXMP2* (peroxisomal membrane protein 2) and *PPIB* (peptidylprolyl isomerase B)(Supplementary Table S3).

### Enrichment in diabetes relevant tissues

We sought to investigate the role of different tissues in the pathogenesis of T2D by looking at the enrichment of significant associations in each tissue. We used the average significance (represented by the squared Z-score averaged across all genes) as a measure of enrichment but recognized the need to account for differential power to detect associations given the different sample sizes used in the training of different tissue models. The enrichment increases as sample size increases (Spearman’s rank correlation *ρ* = 0.887, *p* = 4.69 × 10^−16^) (Figure 3). As expected, we found liver, pancreas, subcutaneous adipose tissue, and skeletal muscle ranked higher than other tissues with similar sample size.

**Figure 3.**
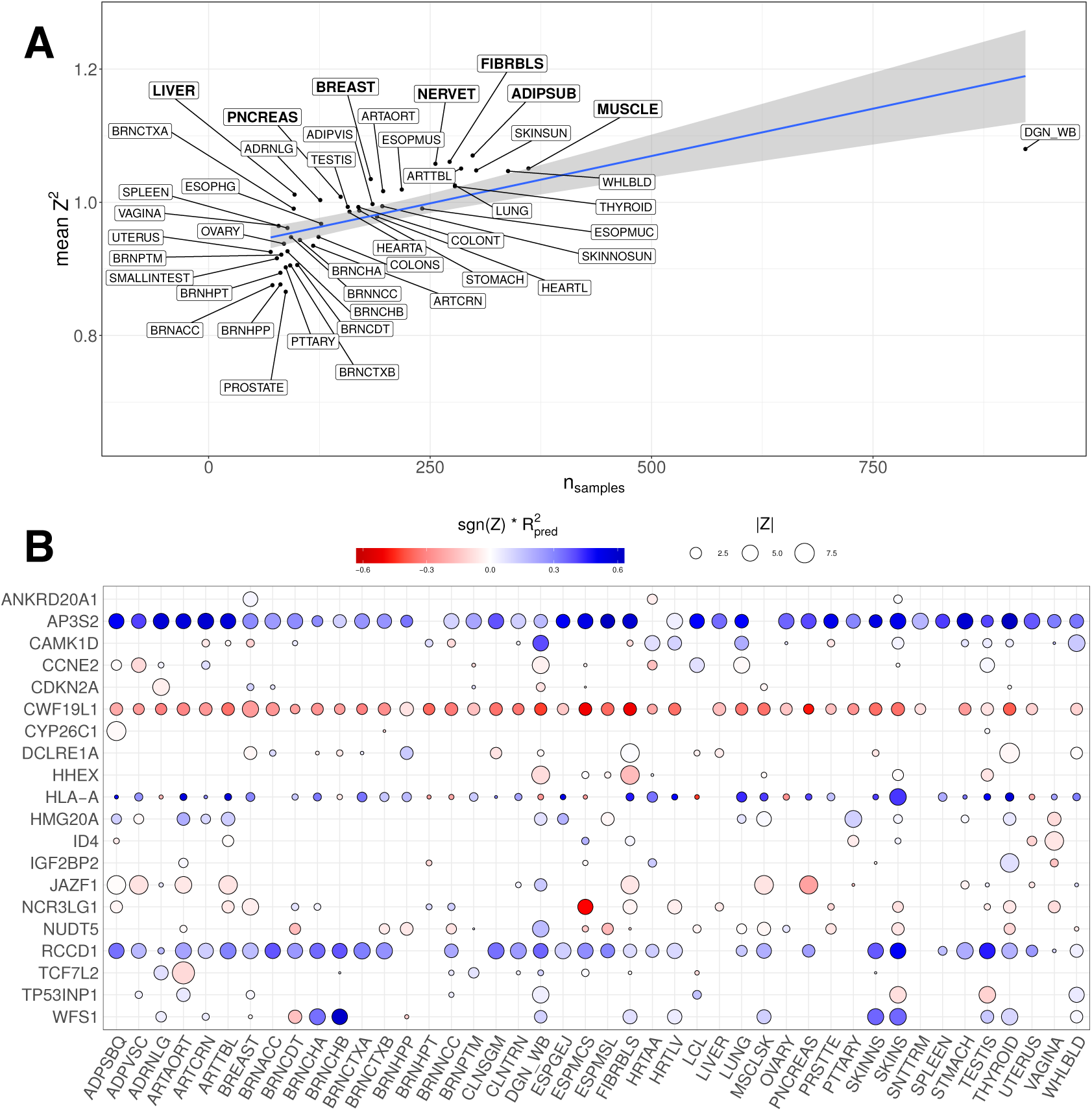
A. Average enrichment of association results vs sample size. Average of Zscore^2^ across all genes is plotted against the number of samples used for the training of the tissue specific models. The enrichment increases with sample size. Reassuringly, diabetes relevant tissues such as liver, pancreas, adipose, and muscle (highlighted) show up at the top of the tissue list for given sample size. **B. Significance of top T2D-associated genes across tissues.** Z-scores of the association between predicted expression levels and T2D case control status across 44 human tissues is shown. Except for *RCCD1, CWF19L1,* and *AP3S2,* genes are associated in only a few tissues indicating context specificity. The size of the circles represent the magnitude of the Z-score. Blue color represents positive association, i.e. increase in expression level associated with increase in T2D risk. Red color represents negative association. The intensity of the color represents the performance of the prediction models (correlation squared between predicted and observed expression levels cross-validated in the training samples). Therefore larger circles indicate more significant associations whereas darker colors indicate higher prediction confidence. Missing circles mean that the association was not performed because of missing model (no good prediction model) or because the prediction SNPs were absent in the GWAS. **Tissue abbreviations**: Adipose - Subcutaneous (ADPSBQ), Adipose - Visceral (Omentum) (ADPVSC), Adrenal Gland (ADRNLG), Artery - Aorta (ARTAORT), Artery - Coronary (ARTCRN), Artery - Tibial (ARTTBL), Bladder (BLDDER), Brain - Amygdala (BRNAMY), Brain - Anterior cingulate cortex (BA24) (BRNACC), Brain - Caudate (basal ganglia) (BRNCDT), Brain - Cerebellar Hemisphere (BRNCHB), Brain - Cerebellum (BRNCHA), Brain - Cortex (BRNCTXA), Brain - Frontal Cortex (BA9) (BRNCTXB), Brain - Hippocampus (BRNHPP), Brain - Hypothalamus (BRNHPT), Brain - Nucleus accumbens (basal ganglia) (BRNNCC), Brain - Putamen (basal ganglia) (BRNPTM), Brain - Spinal cord (cervical c-1) (BRNSPC), Brain - Substantia nigra (BRNSNG), Breast - Mammary Tissue (BREAST), Cells - EBV-transformed lymphocytes (LCL), Cells - Transformed fibroblasts (FIBRBLS), Cervix - Ectocervix (CVXECT), Cervix - Endocervix (CVSEND), Colon - Sigmoid (CLNSGM), Colon - Transverse (CLNTRN), Esophagus - Gastroesophageal Junction (ESPGEJ), Esophagus - Mucosa (ESPMCS), Esophagus - Muscularis (ESPMSL), Fallopian Tube (FLLPNT), Heart - Atrial Appendage (HRTAA), Heart - Left Ventricle (HRTLV), Kidney - Cortex (KDNCTX), Liver (LIVER), Lung (LUNG), Minor Salivary Gland (SLVRYG), Muscle - Skeletal (MSCLSK), Nerve - Tibial (NERVET), Ovary (OVARY), Pancreas (PNCREAS), Pituitary (PTTARY), Prostate (PRSTTE), Skin - Not Sun Exposed (Suprapubic) (SKINNS), Skin - Sun Exposed (Lower leg) (SKINS), Small Intestine - Terminal Ileum (SNTTRM), Spleen (SPLEEN), Stomach (STMACH), Testis (TESTIS), Thyroid (THYROID), Uterus (UTERUS), Vagina (VAGINA), Whole Blood (WHLBLD).

However, when we examined individual genes, many of established T2D genes (e.g. *TCF7L2, WFS1, IRS1*) show associations in tissues that are not traditionally considered relevant for diabetes. For example, *KCNJ11,* which encodes a potassium ion transporter in pancreatic islet, *β*-cells and plays an integral role in glucose-stimulated insulin secretion [25], was significantly associated in esophagus, skin, and whole blood whereas *TCF7L2* association was only detected in aortic artery. *WFS1,* known to cause a syndromic form of diabetes, was significantly associated with T2D in multiple tissues but none of the top tissues (skin, tibial nerve, and thyroid) are among diabetes-relevant ones.

Among the top 20 genes (stringent Bonferroni significant) only three (*RCCD1, CWF19L1,* and *AP3S2*) show significance across many tissues. For the majority of genes, the association is only detected in a handful of tissues (Figure 3 B). This is probably a consequence of the context specificity of regulatory mechanisms that lead to disease in the pathogenic tissue. However, because of sharing of regulatory mechanisms across tissues and because we are examining a large number of tissues, i.e. multiple experiments, we are able to detect the relevant regulatory mechanism, which may or may not be the causal tissue, but happened to have the right environmental or context trigger.

Given the complexity of gene regulation such as context specificity, feedback loops, as well as hidden confounders in the expression data, the regulatory activity may not always be detected in the tissue most relevant to the pathobiology of an implicated gene. But because of sharing of regulation across tissues [9], an agnostic scanning of multiple tissues provides us with additional windows of opportunity to detect the relevant regulatory activity.

### Monogenic diabetes genes enriched in T2D associations

Next we asked if the modest changes in the expression of genes involved in monogenic forms of diabetes could affect the risk of T2D. For this purpose, we examined the enrichment of significant MetaXcan associations among genes involved in monogenic forms of diabetes from [26]. Figure 4 shows the qq-plot for the full set of genes in black, the 81 monogenic diabetes related genes in blue, the smaller list of 28 monogenic genes for which T2D was the primary phenotype. Interestingly, monogenic diabetes related genes (blue) are more significantly associated than others (further away from the gray identity line) and the enrichment increases for genes where diabetes is the primary phenotype (green). Diabetes genes from ClinVar and OMIM showed enrichments in between the two diabetes gene sets.

This result supports the model of a continuum of diabetes phenotypes [27] (from severe to milder forms) in which rare loss of function variants cause severe forms of diabetes whereas smaller alterations of the expression levels of the same genes increase the risk of a later-onset T2D.

**Figure 4.**
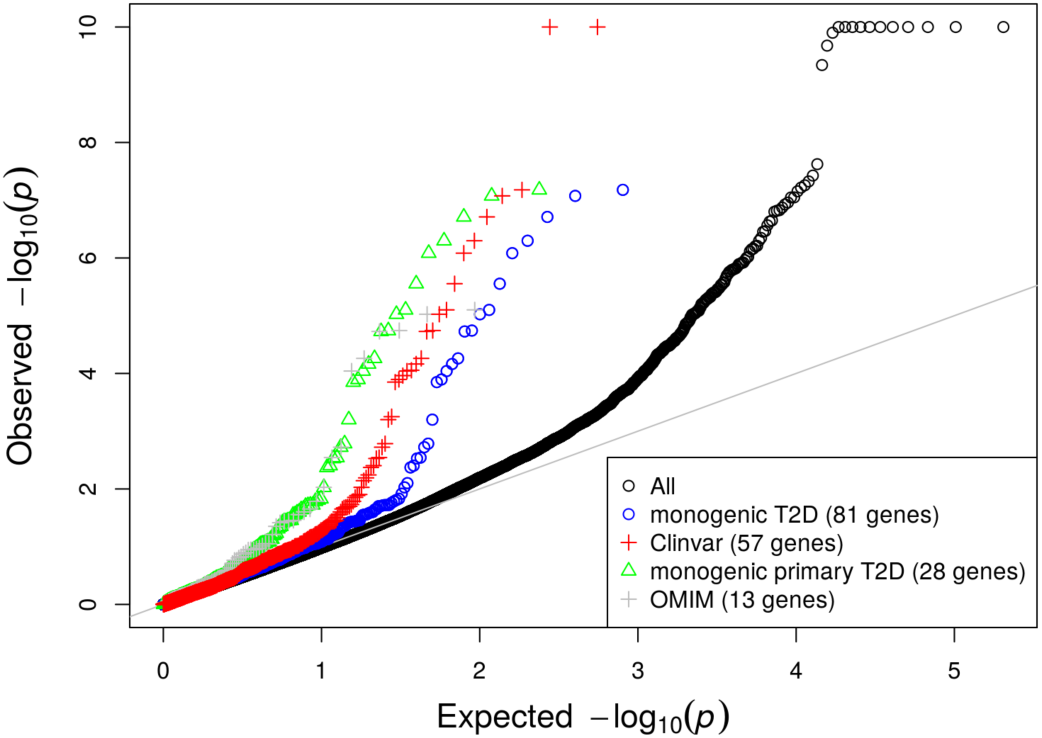
Enrichment of monogenic diabetes genes among T2D associations. This figure shows the qqplot of the p-values of the association between predicted expression levels and T2D case status from the trans-ethnic study across 44 human tissues. The black circles denote the full set of results (204,981 gene-tissue pairs). The blue circles correspond to the qqplot of the monogenic diabetes genes from [26]. Green circles correspond to a subset of the monogenic gene list, considered to cause diabetes as the main phenotype. We see that monogenic genes are enriched in significant genes (away from the gray line) and that the subset where the phenotype is diabetes is even more enriched (further away from the gray identity line). OMIM (gray +) and ClinVar (red +) list yield similar enrichments. p-values below 10^−10^ have been capped to 10^−10^ for better visualization.

### Analysis of known T2D loci prioritizes effector genes

MetaXcan provides a principled way to prioritize effector genes in known trait-associated loci. To implement this, we defined 68 non-overlapping windows comprising known T2D-associated SNPs (see details in Methods) which we refer to as T2D-loci. We profiled these loci according to the strength, number, and proximity of predicted gene expression associations. We used two thresholds for the multiple test correction: one very stringent that accounts for the total number of tissue/gene pairs (genome-wide *p* = 0.05/204, 981 = 2.4 × 10^−7^) and another one more appropriate for a locus-specific analysis that accounts for the total number of tests within the locus *(locus-wide p* threshold varies by locus).

We found that 33 loci had *at least one* locus-wide significant association (Supplementary Table S9). The significance of the associations are depicted for each of the 33 loci in Supplemental Figures S9 and S10. Nine loci show significance only for the reported gene (*BCL11A, IRS1, FHIT, SLIT3, ETV1, STARD10, KLHL42, C2CD4A,* and the three reported genes at the locus spanning *KCNJ11, ABCC8,* and *NCR3LG1*), 14 show both reported and novel genes, and 9 only show novel genes (Supplementary Table S9). We next highlight a few loci of interest.

### JAZF1 locus

At the window comprising T2D-associated SNPs at the *JAZF1* locus, we observed multiple tissue-level associations for *JAZF1,* the reported T2D gene in this region. *Decreased* expression of *JAZF1* in multiple tissues (inlcuding skeletal muscle, adipose, and pancreas) was associated with T2D (Table S1 and Figure 5A). In addition to *JAZF1,* we find that *increased* expression of *HOXA11,* an upstream transcription factor-encoding gene, is associated with T2D at the *locus-wide* level (Figure 5A).

**Figure 5.**
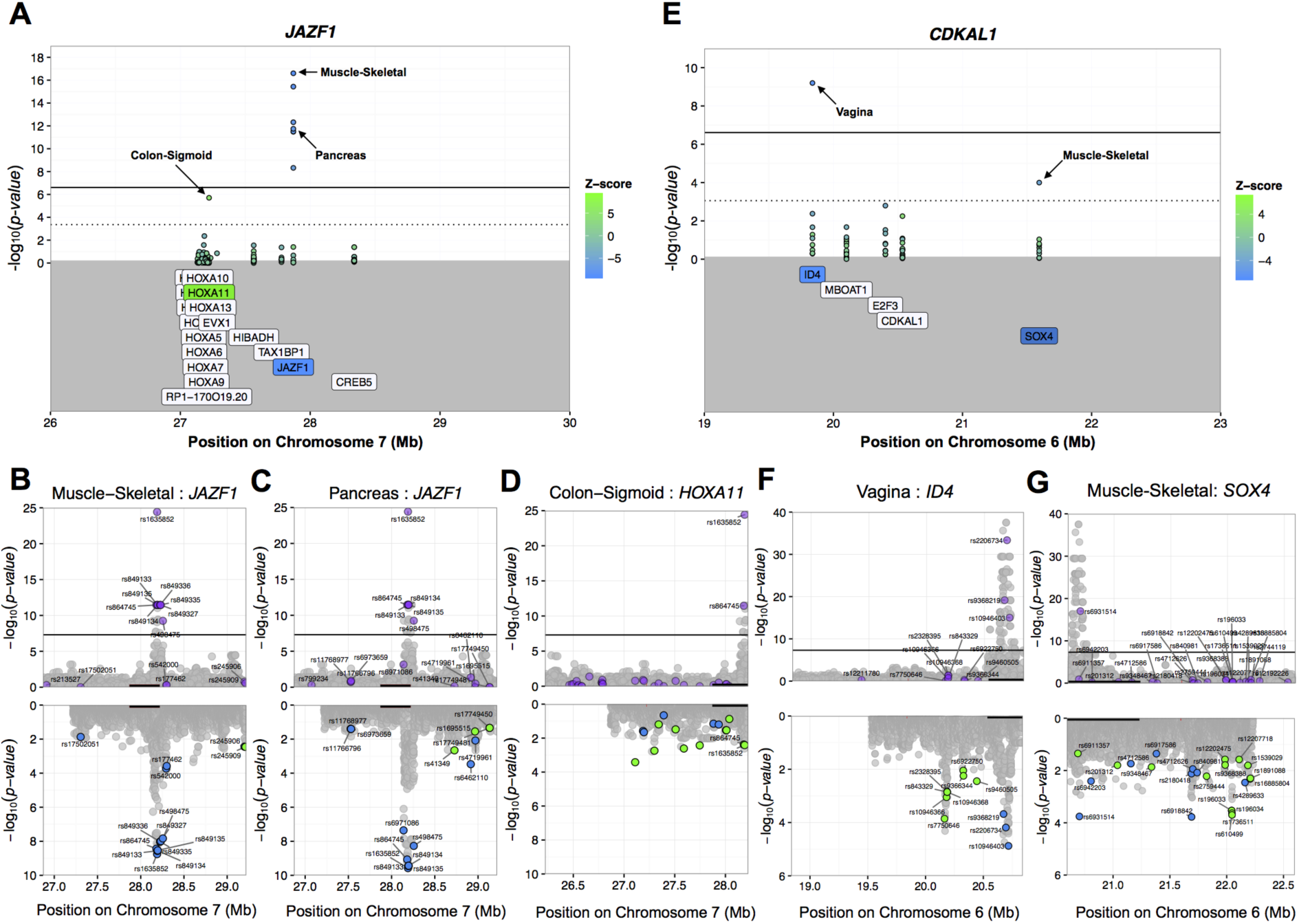
Predicted gene expression analysis identifies novel gene associations implicating distal genes at T2D loci. (A) MetaXcan association plot at the *JAZF1* locus. Solid and dotted lines denote Bonferroni (cross-tissue) and locus-level significance thresholds, respectively. Green and blue fill indicate positive and negative direction of associations (i.e. sign of Z-score), respectively. Label shading shows direction for the top tissue-level association for each meeting MetaXcan significance thresholds. Miami plots showing GWAS (*Upper*) and GTEx V6p eQTL (*Lower*) association p-values for SNPs in the gene expression prediction models for (B) *JAZF1* in skeletal muscle, (C) *JAZF1* in pancreas, and (D) *HOXA11* in sigmoid colon. Color in the eQTL plots indicates direction of effect for the disease-promoting allele of each predictor SNP with green and blue denoting positive and negative effects on gene expression, respectively. Black line segment in each plot shows interval spanned by gene start and end sites. *HOXA11* represents a distal novel T2D gene at the *JAZF1* locus that shares two predictor SNPs (rs1635852 and rs864745) with *JAZF1* in the muscle and pancreas models, respectively. (E) MetaXcan association plot at the *CDKAL1* locus. Miami plot of GWAS and eQTL association profiles for the (F) *ID4* in vagina and (G) *SOX4* in skeletal muscle models. Both *ID4* and *SOX4* represent distal novel T2D genes at the *CDKAL1* locus.

To gain further insight into these associations, we examined the effect of the SNPs that make up the prediction models on the phenotype and on the expression of the corresponding gene. We find that many of the SNPs in the prediction models for *JAZF1* fall within eQTL association peaks in these tissues and are themselves significantly associated with T2D (Figure 5B-C). Moreover, the disease-promoting alleles for these SNPs are associated with decreased expression of *JAZF1* (Figure 5B-C). However, the lead SNP in the *JAZF1* prediction models (rs1635852) is also present in the model for *HOXA11* expression where the disease-promoting allele associates with increased gene expression (Figure 5B-D).

### *CDKAL1* locus

The reported gene, *CDKAL1,* showed no significant association whereas predicted gene expression of nearby genes *ID4* and *SOX4* associated with T2D (Figure 5E). Although there were multiple eQTL association peaks evident among the set of SNPs in the prediction models for these genes, there was only one GWAS peak in this region. Moreover, the disease-promoting alleles of the model SNPs within the shared peak associated with a decrease in gene expression 5F-G.

### *AP3S2, PRC1,* and *VPS33B* locus

We observed the most tissue-level gene associations at a region spanning three *reported* T2D genes: *AP3S2, PRC1,* and *VPS33B* (Figure 6A). Although each of these putative T2D genes were supported by our MetaXcan analysis, the most significant associations at this region corresponded to a *novel* T2D gene,*RCCD1,* with increased expression of this gene associated with T2D(Figure 6A, Table S1, and Supplementary Table S9).

**Figure 6.**
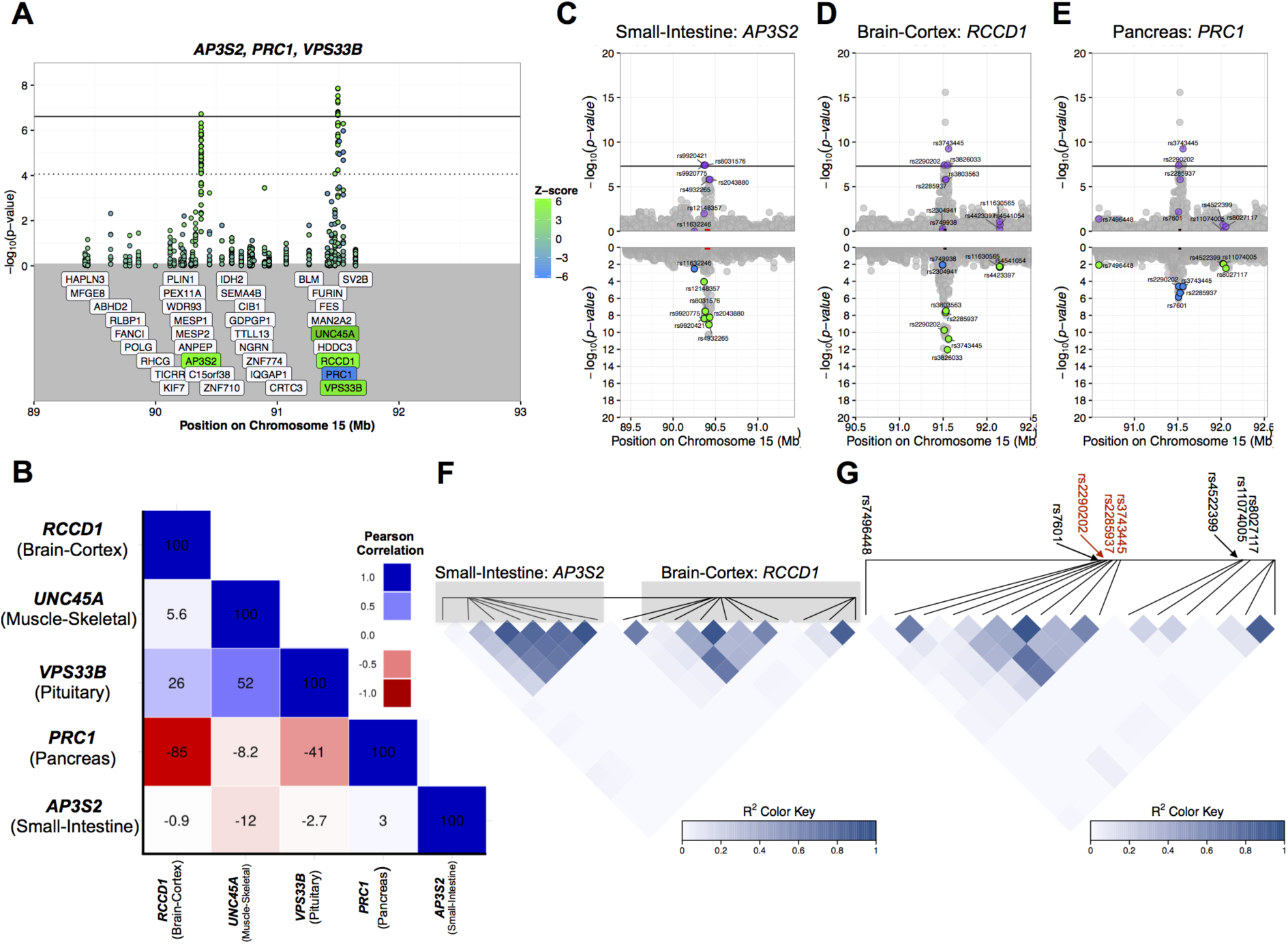
*RCCD1* shows multiple tissue-level associations at a region spanning three *reported* T2D genes. (A) MetaXcan association plot at the region comprising *AP3S2*, *PRC1,* and *VPS33B.* Solid and dotted lines denote Bonferroni (cross-tissue) and locus-level significance thresholds, respectively. Green and blue fill indicate positive and negative direction of associations (i.e. sign of Z-score), respectively. Label shading shows direction for the top tissue-level association for each meeting MetaXcan significance thresholds. (B) Correlation plot of predicted gene expression values in GTEx V6p for the top MetaXcan tissue-level gene associations implicated in the region. Miami plots showing GWAS *(Upper*) and GTEx V6p eQTL (*Lower*) association p-values for SNPs in the gene expression prediction models for (C) *AP3S2* in small intestine, (D) *RCCD1* in brain cortex, and (E) *PRC1* in pancreas. Color in the eQTL plots indicates direction of effect for the disease-promoting allele of each predictor SNP with green and blue denoting positive and negative effects on gene expression, respectively. Black and red line segments in each plot shows interval spanned by *PRC1* and each predicted gene, respectively. (F) LD heatmap of the full set of predictor SNPs in the *AP3S2* (small intestine) and *RCCD1* (brain cortex) models. (G) LD heatmap of the full set of predictor SNPs in the *PRC1* (pancreas) and *RCCD1* (brain cortex) models. All *PRC1* model SNPs are labeled with red color denoted SNPs shared between the two models. *RCCD1* represents a novel gene association with uncorrelated predicted gene expression with *AP3S2* (small intestine) but highly correlated with *PRC1* (pancreas) predicted expression.

The only other gene in this interval supported by at least one genome-wide significant association was the *reported* T2D gene *AP3S2* where increased expression was associated with disease status. Moreover, increased expression of *AP3S2* in 29 tissue models associated with T2D at the window-level threshold (Figure 6A and Supplementary Table S9).

The variants underlying the top associations for *RCCD1* and *AP3S2* are independent from each other as the SNPs constituting the respective predictive models are not in linkage disequilibrium with each other (Figure 6F). The genetically predicted gene expression values for *RCCD1* in brain cortex and *AP3S2* in small intestine are also uncorrelated (Figure 6B). However, the genetically predicted gene expression values for the *RCCD1* in brain cortex and *PRC1* in pancreas (the top model for this *reported,* T2D gene), are strongly and negatively correlated with each other, consistent with their directions of association with T2D (Figure 6A-B). The predictive models underlying these associations share three SNPs in common (rs2290202, rs2285937, and rs3743445) that are associated with increased expression of *RCCD1* in brain cortex and decreased expression of *PRC1* in pancreas (Figure 6D-E,G). Therefore, the top tissue-level gene associations for *RCCD1* and *PRC1* are likely driven by the same regulatory variants with pleiotropic effects on gene expression.

### Strong GWAS signals may act through the regulation of multiple genes

Given the preponderance of loci (21/33) where the predicted expression of multiple adjacent genes associated with T2D (e.g. *RCCD1*, *PRC1, VPS33B*, and *UNC45A* at the *PRC1* locus), we hypothesized that stronger SNP associations from GWAS involve SNPs with pleiotropic effects on gene expression. Indeed, we observed a correlation between the strength of the top T2D-associated SNP within a genomic region and the number of MetaXcan-implicated genes (Spearman’s *ρ* = 0.43, *p* = 2.8 × 10^−4^) (Figure 7A). At the region spanning *TCF7L2,* tissue-level associations implicate 6 genes, including *TCF7L2* itself. Decreased expression of *TCF7L2* in aortic artery and increased expression in thyroid associated with T2D and genome-wide at window-level significance, respectively (Figure 7B). However, we found that the genetically predicted gene expression values corresponding to all tissue-level gene associations at window-level significance were correlated with that for *TCF7L2* in aortic artery in directions consistent with the directions observed in the association plot (Figure 7B-C). Moreover, these tissue-level gene associations likely share a regulatory genetic basis as SNPs across predictive models fall within with same GWAS association peak and are in linkage disequilibrium with predictor SNPs for *TCF7L2* in aortic artery (Figure 7D-G)

**Figure 7.**
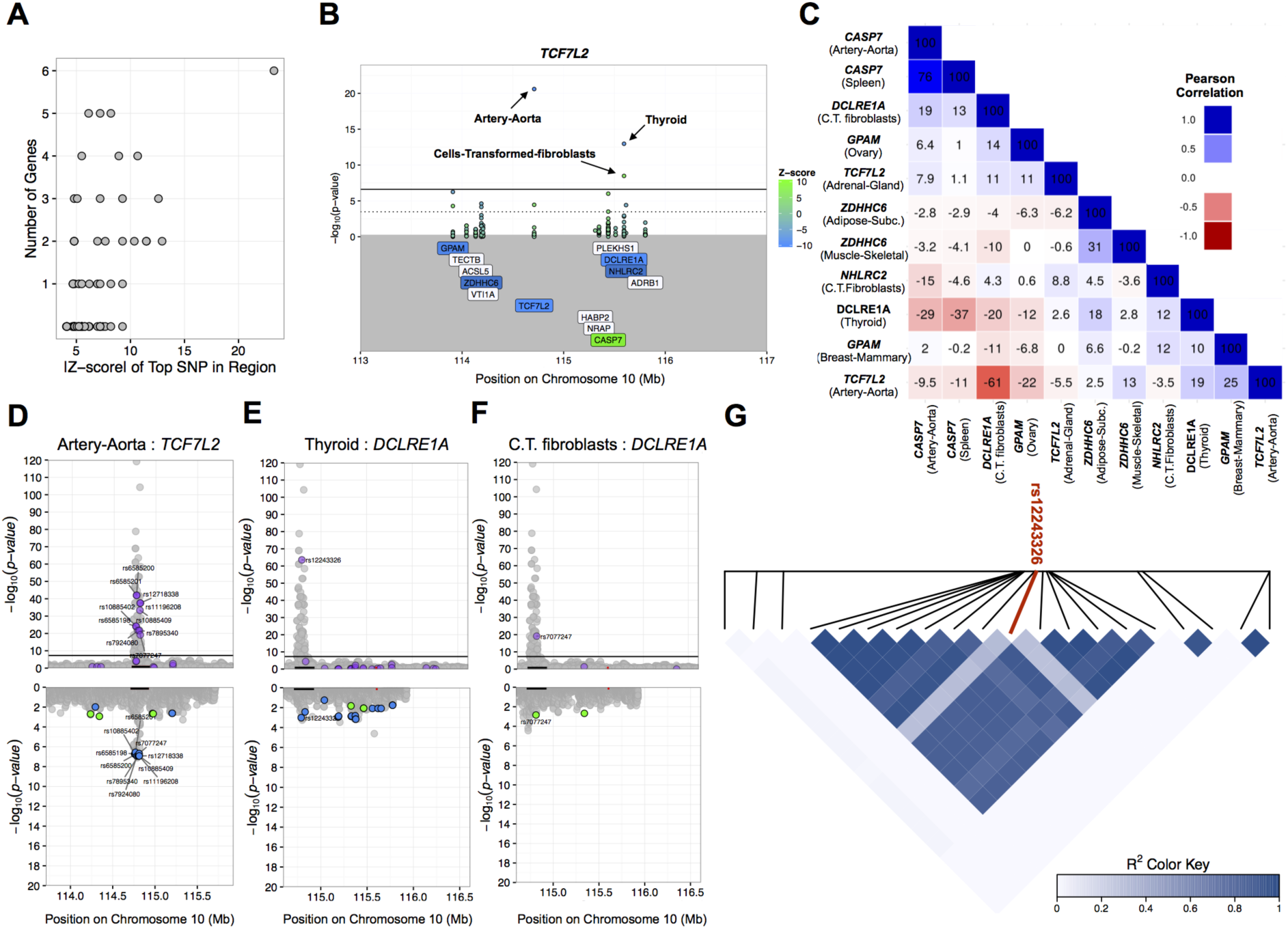
Locus analysis identifies multiple correlated gene associations at the *TCF7L2* locus. (A) Absolute value of Z-score for the top T2D-associated SNP in each non-overlapping region is shown against the number of genes implicated by MetaXcan in each region (*p* ≤ locus threshold). (B) MetaXcan association plot at the *TCF7L2* locus. Solid and dotted lines denote Bonferroni (cross-tissue) and locus-level significance thresholds, respectively. Green and blue fill indicate positive and negative direction of associations (i.e. sign of Z-score), respectively. Label shading shows direction for the top tissue-level association for each meeting MetaXcan significance thresholds. Tissue models are indicated for each of the three gene associations meeting the cross-tissue, genome-wide Bonferroni threshold. (C) Correlation plot of predicted gene expression values in GTEx V6p for the MetaXcan tissue-level gene associations meeting locus-level significance in the region. Miami plots showing GWAS (*Upper*) and GTEx V6p eQTL (*Lower*) association p-values for SNPs in the gene expression prediction models for (C) *TCF7L2* in aortic artery, (D) *DCLRE1A* in thyroid, and (E) *DCLRE1A* in transformed fibroblast cell lines. Color in the eQTL plots indicates direction of effect for the disease-promoting allele of each predictor SNP with green and blue denoting positive and negative effects on gene expression, respectively. Black and red line segments in each plot shows interval spanned by *TCF7L2* and *DCLRE1A,* respectively. (G) LD heatmap of prediction model SNPs for *TCF7L2* (aortic artery) and top SNP in prediction model for *DCLRE1A* in thyroid (red), rs12243326.

## Discussion

We performed a large-scale, *in silico* study of predicted gene expression across a comprehensive set of human tissues to prioritize genes that alter the risk of T2D through regulation of gene expression levels. We corroborated many of the known T2D genes, which supports the role of regulation of gene expression levels as a key mediating mechanism, but also found novel genes both in known loci as well as in completely novel loci. Replication in an independent cohort gives further support to our results.

Among novel genes of interest are genes previously linked to related traits such as *HKDC1* (pregnancy-related glycemic traits) and *GPAM* (LDL cholesterol). Moreover, pathogenic variants in *KCNK17* - a novel T2D gene implicated by MetaXcan - have been identified in patients with hyperinsulinemic hypoglycemia and cardiac arrhythmia [28]. Another example, *SOX4,* has been implicated with diabetes in multiple experiments. The expression of *SOX4*/Sox4b has been shown to play a role in pancreas development and insulin secretion in mouse models and human cell lines [29, 30, 31]. For example, mice expressing a mutant form of Sox4 exhibited a 40% reduction in glucose-induced insulin secretion [32]. Moreover, overexpression of *SOX4* in a human insulin-secreting cell line (Endo-C-βH2) resulted in a marked decrease in insulin release through up-regulation of *STXBP6* - a gene encoding an exocytosis-regulating protein [32].

Averaging across the genome, we found that diabetes relevant tissues such as liver, pancreas, adipose, and muscle are enriched with significant associations. However, when we look into individual genes at the top significance level we found associations in tissues that are not typically linked to diabetes. For example, *TCF7L2* was significantly associated only in aortic artery and adrenal gland, which is a consequence of the fact that with GTEx samples active regulation of this gene was only found in these tissues.

Most associations were discovered in a few tissues indicating strong context specificity. Some of the associations may be pointing to a real causal tissue but others are likely to be a consequence of shared regulation across tissues. Although the context specificity limits our ability to detect associations even in causal tissues, the sharing of the regulation across multiple tissues opens additional opportunities for discovering the disease-causing regulatory mechanism albeit in non-causal tissues where the environmental conditions were met. Our results underscore the benefits of an agnostic scanning across all available tissue models.

An important caveat of this study is that we used average expression over a gene when generating predictive models and may therefore miss the consequences of regulatory variants that impact splicing at T2D loci. Although it should not create false positives, this may explain why we failed to detect genome-wide significant associations at some regions encompassing putative T2D genes.

The predictive models employed in this study were trained from local variants within 1 Mb of each gene. Although most eQTLs mapped in human tissues are local eQTLs, this is influenced by the fact that the greater number of genetic variants, smaller haplotype structure, and relative smaller sample sizes associated with human studies considerably reduces power to detect distal eQTLs that regulate target genes through a non allele-specific mechanism (i.e. *trans* eQTLs) [7]. However, distal-acting eQTLs mapped in pancreatic islet and insulin-responsive peripheral tissues may account for some of the genetic architecture of T2D [33, 16].

In our study, we applied MetaXcan to explicitly integrate regulatory genetic information to improve disease gene mapping and overcome key limitations of GWAS and differential gene expression studies [21]. This approach, along with similar approaches adopted by Gusev *et a1*. (2015) and Zhu *et a1*.(2016), directly addresses the importance of eQTLs in complex human traits and advances genetic studies beyond GWAS [34, 35]. Importantly, we provide information about the direction of gene expression that associates with disease, that was predominantly consistent across the most significant associations discovered in this study and replicated in an independent cohort. This immediately suggests potential therapeutic targets where the increased expression of genes - many of which were not previously reported from GWAS - significantly relates to increased disease risk. Moreover, these results establish a basis for subsequent experiments (e.g. gene editing) to interrogate the cellular and physiological consequences of dysregulation of novel candidate genes. Therefore, this investigation represents an important step forward in elucidating the genetic basis of T2D and other complex diseases.

## Materials and Methods

No identifiable data were used for this study, which was considered to be “Non human subject research”.

### Determining SNP predictors of gene expression

#### DGN whole blood model

SNP predictors of gene expression in whole blood tissue were determined as described in [36] with genome-wide genotype and RNA-seq data from the Depression Genes and Networks (DGN) cohort study [10] corresponding to 922 unrelated individuals (π̂ < 0.05) of European ancestry. In brief, imputation of 650K SNPs with minor allele frequency (MAF) > 0.05 and non-significant departure from Hardy-Weinberg equilibrium (HWE) were imputed to a 1000 Genomes (Phase 1, version 3) reference panel [37] with ShapeIt2 [38]. The full set of ~1.9 M imputed SNPs with MAF > 0.05 and imputation *R*^2^ > 0.8 were subsetted to SNPs included in HapMap Phase II [39]. HCP (hidden covariates with prior) normalized gene-level expression data was downloaded from the NIMH repository [36].

#### GTEx tissue models

RNA-seq gene expression from 8, 555 tissue samples (representing 53 unique tissue types) from 544 subjects and imputed genotypes (available for 450 subjects and imputed to a 1000 Genomes reference panel) was obtained from the Genotype Tissue Expression Project (GTEx) data release on 2014-06-13 [9]. Expression measures from the top 44 GTEx tissues with the largest available sample sizes [9, 36] and SNPs included in HapMap Phase II (~ 2.6 M) were carried forward in our model fitting procedure.

In order to delineate a set of informative SNPs for predicting tissue-level gene expression, we performed penalized regression with the Elastic Net - a multivariate linear model that includes the *l*_2_-norm and *l*_1_*-* norm penalties from ridge regression and the Least Absolute Shrinkage and Selection Operator (LASSO) procedure, respectively [40, 41]. This method leverages shrinkage parameters that enable feature selection while solving for the coefficient solutions to the regression of gene expression on SNP genotypes. The Elastic Net model includes an additional mixing parameter a that determines the contribution from each penalty parameter (i.e. the Elastic Net model is equivalent to ridge regression and LASSO regression when α = 0 and α =1, respectively) [41]. Here, we set α = 0.5. Gene expression - as measured by reads per kilobase of transcript per million reads mapped (RPKM) - was adjusted for potential batch effects and unmeasured confounders by regressing out the first 15 PEER factors [42] in R [43]. For each gene expressed in a tissue, model fitting was performed by regressing PEER-adjusted gene expression on the set of SNPs located within 1 Mb of the transcription start site (TSS). Therefore, subsequent analyses pertain to estimates of genetic components of gene expression attributable to local regulatory variants. The SNP coefficients from this procedure are used as weights to estimate the genetic component of gene expression and are publicly available (http://predictdb.org).

### Summary data from GWAS on type 2 diabetes

#### Trans-ethnic Study summary data

Input GWAS summary data used in our MetaXcan-based association of predicted gene expression and T2D corresponded to the trans-ethnic meta-analysis study [4] and was publicly available and downloaded from the DIAGRAM Consortium website (http://diagram-consortium.org/). This study involved a meta-analysis of 26, 488 cases and 83, 964 controls subjects from populations of European, east Asian, south Asian, and Mexican, and Mexican American ancestry. Although a majority of individuals were of European ancestry (12,171 cases and 56, 862 controls) [44], the study included East Asian individuals from the AGEN-T2D Consortium (6, 952 cases and 11, 865 controls) [45], south Asian individuals from the SAT2D Consortium (5, 561 cases and 14, 458 controls) [46], and individuals of Mexican and Mexican American ancestry (1,804 cases and 779 controls) [47]. SNPs were lifted to NCBI build GRCh37 (UCSC hg19 assembly).

#### GWAS on T2D results from GERA study

Replication analyses were performed using summary GWAS data from an analysis on the Genetic Epidemiology on Adult Health and Aging (GERA) cohort (dbGaP phs000674.p1). GERA represents a large, multi-ethnic cohort of individuals of European, East Asian, African American, and Latino ancestry where each subgroup was genome-wide genotyped with arrays designed to maximize coverage of common and low-frequency variants in each constituent population [48, 49]. T2D case status was determined from ICD-9 codes available from electronic medical health records. SNPs meeting selection criteria for MAF (≥ 1%), HWE departure (*p* > 10^−6^), and call rate (> 95%) were pre-phased with SHAPEITv2.5 [38] and imputed to a 1000 Genomes reference panel with IMPUTEv2.3 [50]. GWAS on T2D was performed on a set of 71, 604 unrelated (π̂< 0.2) subjects (9, 747 cases and 61, 857 controls) with SNPTESTv2.5 [51] and adjusted for principal components (PCs) to correct for population stratification.

#### Gaussian Z-score imputation of GWAS summary statistics

There was a total of 1, 803, 748 SNPs in the gene expression prediction models across all 44 tissue models (including whole blood from the DGN study) that resulted from our Elastic Net fitting procedure (*α* = 0.5) and corresponded to genes with prediction FDR < 0.05. However, not all of these SNPs were present in the summary data from the trans-ethnic and GERA meta-analysis of GWAS - the coverage of models SNPs in these datasets was 93.4% and 71.3%, respectively. In order to improve coverage of *model* SNPs to further enable comparisons between these summary datasets in replication and meta-analyses of MetaXcan results, we applied Gaussian Z-score imputation of GWAS summary statistics as implemented in Imp-G Summary software (http://bogdan.bioinformatics.ucla.edu/software/) [52]. We imputed GWAS Z-scores to a reference panel of all available ancestral populations from the 1000 Genomes Project phase 1 (v3) <u>r</u>elease [37]. The requisite haplotype files in Beagle [53] format were accessed from http://bochet.gcc.biostat.washington.edu/beagle/1000_Genomes.phase1_release_v3/ on August 1, 2016. We restricted imputed *model* SNPs to those with imputation quality score (*R*^2^-pred) ≥ 0.80 [52]. This increased coverage of *model* SNPs in the trans-ethnic and GERA GWAS summary datasets to 96.1% and 91.6%, respectively.

### Testing for association between predicted gene expression and T2D with MetaXcan

For this study, we used MetaXcan [22], an extension of the PrediXcan method [21], that takes as input summary statistics from GWAS. This approach improves computational efficiency over PrediXcan as it does not require individual-level genotype data to estimate genetic components of gene expression for subsequent trait association testing. Rather, the PrediXcan Z-statistic (*Z_g_*) is approximated by:

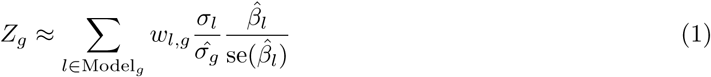

where *w_l_*, _*g*_ represents the prediction model weight for SNP *l* on gene *g*, *σ_l_* is the standard deviation for SNP *l*, *σ̂_g_* is the standard deviation of predicted expression for gene *g*, *β̂_l_* is the regression coefficient for the regression of expression on the allelic dosage of SNP *l*.

In our MetaXcan analyses of T2D, we use regression coefficients (*β̂_l_*) from results from the trans-ethnic meta-analysis of GWAS and the GWAS on T2D from the GERA study. Values for *w_l,g_* were generated as described above and available from the PredictDB website (http://predictdb.org). *σ̂_g_*^2^ is estimated as:

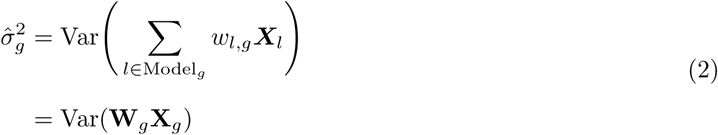

Where **W**_*g*_ is the vector of *w_l,g_* for SNPs in the model of *g* and Var(**X**_*g*_) is the covariance matrix of **X**_*g*_. We use SNP information from a 1000 Genome Project reference panel (European ancestry) to the compute the variances and covariances of the SNPs used to predict gene expression.

### Locus analysis of T2D-associated regions

We first identified a set of 111 reported T2D genes based on their being the most proximal to T2D-associated SNPs at *p* < 5 × 10^−6^ in the trans-ethnic meta-analysis of GWAS. We then delineated genomic regions for each reported gene by taking the set of all significantly-associated SNPs annotated to that gene and demarcating a window bounded by the SNPs most distal to each other. We then expanded the region by 1 Mb upstream and downstream of the “boundary” SNPs. This ensured that the reported gene was included within the genomic window corresponding to the T2D-associated locus. This procedure resulted in 68 non-overlapping genomic regions (i.e. windows).

We then performed a genome-wide MetaXcan analysis of the trans-ethnic study to test for association between predicted expression for each gene with prediction FDR ≤ 0.05 (from the regression of observed gene expression on predicted gene expression) in each of the 44 tissue models described above. When visualizing the MetaXcan results at each T2D locus we considered two significance thresholds: (1) genome-wide significance correcting for the total number of tests performed across all available tissue models and significance correcting for the number of tests performed within each non-overlapping region.

### Meta-analysis of association results from the MetaXcan analysis of trans-ethnic and GERA cohorts

We performed a sample-sized based meta-analysis [54] of the association results from our MetaXcan analyses of the trans-ethnic and GERA studies where the Z-score (*Z*) was given by:

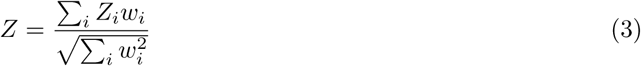

where *Z_i_* = Φ^−1^(*P_i_*|2) * sign(Δ_*i*_), *P_i_* is the p-value for study *i, w_i_* = √(*N_i_*), *N_i_* refers to the sample size for study *i*, Δ_*i*_ is the direction of effect in study *i*, and the overall P-value is given by:

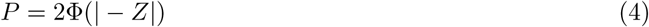

### Gene set enrichment analysis of MetaXcan-significant gene sets

**Gene set enrichment analysis.** Gene set enrichment analyses (GSEAs) were performed by comparing sets of significant genes implicated by our MetaXcan analyses with the complement set of GENCODE v18 [55] genes (~18K) for which we can predict in any tissue model with prediction FDR ≤ 0.05. We restricted analyses to test for enrichment of pathways designated as Gene Ontology Biological Process (GO:BP) [56]. Overrepresented p-values were obtained from a parametric Fishers exact test using the Wallenius approximation and a non-central hypergeometric distribution [57]. GSEA was performed with the **GOseq** package [57] in R [43] that applies a weighting scheme to control for selection bias introduced by differences in transcript length.

### Cross-phenotype comparison of T2D gene enrichment

The full set of annotated single variant results from published GWAS listed on the National Human Genome Research Institute / European Bioinformatics Institute (NHGRI-EBI) online catalogue - corresponding to 1, 362 phenotypes - was downloaded from https://www.ebi.ac.uk/gwas/ (Accessed April 2016). The set of reported genes for each trait was tested for enrichment of genes significantly associated with T2D in our MetaXcan analyses through a resampling procedure. An empirical p-value was determined by first taking the observed count of intersecting genes between reported genes for each trait and MetaXcan-significant T2D genes. We then generated a null distribution of counts by randomly sampling 10,000 gene sets from the set of all GENCODE v18 [55] with prediction FDR ≤ 0.05 in at least one tissue model. Each sample was matched for the number putative genes reported for each trait and the overlap with the set of MetaXcan-significant genes was recorded. The enrichment p-value was calculated as the number of instances a sampled count value equaled or exceeded the observed count between reported trait genes and MetaXcan-significant genes.

## Acknowledgments

This project was funded in part by the Genotype-Tissue Expression project (GTEx) (R01 MH101820 and R01 MH090937), JMT (F31 DK 101202), HKI (R01 MH107666), and the University of Chicago Diabetes Research and Training Center (P30 DK020595). Andrew P Morris is a Wellcome Trust Senior Fellow in Basic Biomedical Science under award WT098017.

## Data Access

### Trans-ethnic Type 2 Diabetes Study dataset

We downloaded trans-ethnic SNP level meta analysis results from http://diagram-consortium.org

### GERA dataset: dbGaP accession phs000674.v2.p2.

Data came from a grant, the Resource for Genetic Epidemiology Research in Adult Health and Aging (RC2 AG033067; Schaefer and Risch, PIs) awarded to the Kaiser Permanente Research Program on Genes, Environment, and Health (RPGEH) and the UCSF Institute for Human Genetics. The RPGEH was supported by grants from the Robert Wood Johnson Foundation, the Wayne and Gladys Valley Foundation, the Ellison Medical Foundation, Kaiser Permanente Northern California, and the Kaiser Permanente National and Northern California Community Benefit Programs. The RPGEH and the Resource for Genetic Epidemiology Research in Adult Health and Aging are described in the following publication, Schaefer C, et al., The Kaiser Permanente Research Program on Genes, Environment and Health: Development of a Research Resource in a Multi-Ethnic Health Plan with Electronic Medical Records, In preparation, 2013.

## Software and code

All code available in https://github.com/hakyimlab/MetaXcan and https://github.com/hakyimlab/MetaXcanT2D

